# Adipocyte-related genes analysis contributes to the prognosis of atrial fibrillation

**DOI:** 10.64898/2025.12.18.695292

**Authors:** Feixing Li, Linbing Liu, Aiai Zhang, Wei Liu, Mengyang Yi, Wei Han, Jianying Song, Yue Shi, Xiaoyuan Wang, Meiling Du, Fangjiang Li

**Author notes:** These authors contributed equally to this work (Feixing Li, Linbing Liu). Corresponding author: Fangjiang Li, Email addresses.

## Abstract

**Background:** Patients with mild left atrial appendage fibrosis have a significantly lower rate of atrial fibrillation (AF) recurrence after 5 years compared to those with moderate or severe fibrosis, which presented a significantly increased positive area of cellular senescence.

**Methods:** GSE135445, a RNA-sequencing dataset between epicardial adipose tissue (EAT) from patients with persistent non-valvular AF and sinus rhythm (SR), was downloaded and analyzed and hub genes were obtained. To obtain the abundance of immune cells, the CIBERSORT algorithm with 100 permutations was applied. The senescent hub genes were obtained and validated using human blood samples.

**Results:** A total of 164 up-regulated and 317 down-regulated DEGs was obtained between EAT from subjects with AF and SR. Using immune infiltration analysis in whole PAT related DEGs, the abundance of M0 macrophages was significantly different between PAT and SAT and 22 M0 macrophages-related DEGs were also obtained, such as MMP7, APOE, CCND1, MMP12 and APOC1. Venn diagram was utilized to obtain the same transcripts between senescence datasets and DEGs and PPI network was used to 94 overlapping genes. The top15-degree senescence adipicytes hub genes were explored. To investigate senescence-related hub genes effects on AF progression, we collected the blood samples of patients with paroxysmal atrial fibrillation and persistent atrial fibrillation. There were no significant differences at baseline characteristics and laboratory examinations at 1d after admission except heart failure history and function examinations, for instance, ALT, LDL, LVEF and LA diameters. Using univariate analysis and multivariate analysis, heart failure history, ALT and LA diameters can be prognostic factors for AF progression. BMP7, COL4A3, COL4A4, CHRD, IRX1, SPON1, ALPL, TCF19 and SIX2 were validated to be highly expressed in persistent atrial fibrillation compared to paroxysmal atrial fibrillation.

**Conclusion:** Based on our current study, BMP7, COL4A3, COL4A4, CHRD, IRX1, SPON1, ALPL, TCF19 and SIX2 were validated to be highly expressed in persistent atrial fibrillation compared to paroxysmal atrial fibrillation, which may be potential therapeutic targets to treat AF progression and improve patients’ prognosis.

## 1. Introduction

Atrial fibrillation (AF) is a common arrhythmia with over 33 million people affected all over the world. The degree of fibrosis in the left atrial wall is a significant predictor of the recurrence of atrial fibrillation, especially in patients who have undergone MRI-guided fibrosis ablation treatment[1]. There are differences in LA remodeling parameters (eg. volume, emptying fraction and strain indices) between patients with and without AF recurrence after ablation[2]. Patients with mild left atrial appendage (LAA) fibrosis have a significantly lower rate of AF recurrence after 5 years compared to those with moderate or severe fibrosis[3].

Left atrial epicardial adipose tissue (EAT) is correlated with obesity (BMI), left atrial volume, and fibrosis. There is no co-localization between EAT regions and fibrotic areas, suggesting that the association might be due to systemic or paracrine mechanisms rather than infiltration of EAT into fibrotic regions[4]. The increased levels of plasma proteins were associated with a heightened risk of all-cause mortality in the HF population, irrespective of the presence or absence of co-existing AF[5]. IMD1-53 reduces the duration and inducibility of atrial fibrillation and atrial fibrosis by inhibiting TGF-β1/Smad3-related fibrosis and TGF-β1/Nox4 activity[6].

Compared with sinus rhythm, left atrial appendages (LAAs) with AF presented a significantly increased positive area of cellular senescence, with upregulated expression of p16, p21 and p53. Suppression of p21 by siRNA reduced TP-induced cell senescence IL-1β, and IL-6 elevation, and partly changed SR-related proteins expression. Moreover, we show that the level of γH2AX, a marker of DNA damage, was higher in AF patients than in sinus rhythm controls[7]. Cellular senescence is a state of growth arrest that occurs after cells encounter various stresses. Senescent cells have the characteristic of secreting inflammatory factors and other bioactive substances, and these secretions ultimately lead to inflammation and tissue fibrosis[8].

The stepwise increase in senescence, prothrombotic, and pro-remodeling markers observed from SR to paroxysmal atrial fibrillation (PAF) to permanent atrial fibrillation (PmAF) points towards a possible network that links the human atrium, senescence burden, endothelial dysfunction, thrombogenicity, and atrial remodeling [9].

In this study, AF-related differential expressed genes (DEGs) were obtained from GSE128188, which contained the RNA-sequencing data of left and right atrial appendages from 5 patients in sinus rhythm (SR) and 5 patients in AF. A protein-protein interaction (PPI) network was carried out to explore the hub genes. To validate the hub genes, GSE62203 about human iPS-derived cardiomyocyte data exposed to glucose, endothelin-1 and cortisol and human blood samples were utilized. AF human blood samples were also collected to investigate the effects of the screened genes on cardiac functions.

## 2. Methods

### 2.1 Microarray data and data processing

Using the keywords “adipose tissue” in “atrial fibrillation”, GSE135445 from the Gene Expression Omnibus (GEO) database was downloaded. The RNA-sequencing between epicardial adipose tissue (EAT) from patients with persistent non-valvular AF (n=6) and sinus rhythm (n=6) were based on Illumina HiSeq 4000 (Homo sapiens). The DEGs were explored using “DESeq2” (version 1.28.1) package in R and an adjusted P value < 0.05 and Log2|FC (Fold Change) | >1 were considered as the cut-off criteria.

### 2.2 Enrichment pathways analysis

GO/KEGG pathway analysis were used to explore the functions of DEGs above mentioned was applied utilizing Xiantao Database (www.xiantao.love) . An adjusted P value < 0.05 were considered as significant.

### 2.3 PPI and the hub genes

To investigate the hub genes, STRING database (https://string-db.org) was used with a combined score >0.4 and the nodes were analyzed using Cytoscape v.3.7.1 . The hub genes were obtained utilizing the Cytoscape plug-in Cytohubba.

### 2.4 Immune infiltration analysis

CIBERSORT is a deconvolution approach to characterize the cell compositions in bulk tissues. To obtain the abundance of immune cells, the CIBERSORT algorithm with 100 permutations was applied, utilizing the LM22 matrix as reference. CIBERSORT outputs a deconvolution p-value for each sample to determine the reliability of the results. In this study, we retained the samples with p < 0.05 to analyze the fractions of immune cells. Then, Pearson correlation analysis was applied to obtain the genes associated with the abundance of immune cells. The DEGs with Pearson correlation coefficients (PCC) >0.4 were regarded as immune cells-related DEGs.

### 2.5 Senescence-related DEGs in epicardial adipose tissues

To investigate the senescence-related DEGs in EAT, GSE35957 were analyzed and an adjusted P value < 0.05 and Log2|FC (Fold Change) | >1 were considered as the cut-off criteria. Venn diagram was utilized to obtain the same transcripts between senescence-related DEGs and AF-EAT-related DEGs. GO/KEGG pathways analysis and PPI network were utilized to obtain the hub senescence-related DEGs in epicardial adipose tissues.

### 2.6 The hub genes and their interactions

The hub genes and their interactions were analyzed using NetworkAnalyst 3.0. Specifically, transcription factor (TF) and gene target data were derived from the ENCODE ChIP-seq data (peak intensity signal <500 and the predicted regulatory potential score <1). miRNA interactions with the screened hub genes were shown using miRTarBase v8.0. The hub DEMRGs-chemicals interactions were shown based on data from the Comparative Toxicogenomics Database (CTD) . The hub DEMRGs-drugs interactions were shown using the DrugBank database (Version 5.0).

### 2.7 The central-marginal dispersion in AF EAT

To explore the central-marginal dispersion in EAT from patients with AF, GSE154436 was downloaded and analyzed using Xiaotao website tool, which included RNA sequencing data about central EAT and marginal EAT from patients with AF. Venn diagram was utilized to obtain the same transcripts between AF-related DEGs and central-marginal dispersion-related DEGs.

### 2.8 Validation using human samples

To further validate the hub genes effects on cardiac functions, the AF patients’ blood samples were collected. The inclusion criteria were as follows: (1) Age ≥18 years old; (2) Symptomatic paroxysmal AF, persistent atrial fibrillation, long-term persistent atrial fibrillation; (3) failure to respond to treatment with one or more antiarrhythmic drugs (such as class IC or Class III antiarrhythmic drugs); (4) discontinue antiarrhythmic drugs for at least 5 half-lives before surgery; (5) The patient is willing to accept RFCA treatment and postoperative follow-up, and agrees to sign informed consent. The exclusion criteria were as follows: (1) Prior ablation of atrial fibrillation; (2) Patients with thrombocytopenia (PLT < 80 × 109/L) or with anticoagulation contraindications (such as warfarin, heparin, direct factor Xa, factor IIa inhibitors, etc.); (3) Left atrial (left auricle) thrombus (by TEE or MSCT examination); (4) Abnormal thyroid function (such as uncontrolled hyperthyroidism); (5) severe liver and kidney dysfunction (AST or AL≥ 3 times the normal upper limit; SCr>3.5mg/dl or Ccr<30 ml/min); (6) Women during pregnancy. Before participating in the study, all individuals provided written informed consent, and the protocols were approved by the First Affiliated Hospital of Hebei North University. The hub genes were determined using a human, respectively. Clinical information and laboratory examinations were also collected. qPCR in different groups was performed using TRIzol and TB Green (TaKaRa, RR820). β -actin was the reference gene, and the 2-ΔΔCt method was utilized.

### 2.9 Statistical analysis

Data are presented as mean ± standard deviation, median (Q1–Q3), or frequency (percentage). Statistical analyses were performed using SPSS 23.0. The Shapiro–Wilk normality test and Weltch’s t-test (two groups) were used and the Spearman correlation analysis was applied between the clinical information and laboratory examinations and protein levels of the screened hub genes. Correlation analysis was also performed to validate the effects of the hub genes on patients with AF, especially on those diabetes patients. Statistically significance was set as p < 0.05.

## 3. Results

### 3.1 Identification of DEGs in EAT in AF and further analysis

A total of 164 up-regulated and 317 down-regulated DEGs was obtained between EAT from subjects with AF and SR, which were mainly involved in collagen-containing extracellular matrix, Cytokine-cytokine receptor interaction, and embryonic skeletal system morphogenesis (Fig. 1A, B; Fig. S1; Tab. 1). To investigate the hub genes of EAT from patients with AF, STRING and Cytoscape plugin Cytohubba were utilized. The top30-degree hub genes between EAT from subjects with AF and SR were screened, including LYVE1, CSF1R, BMP7, CD68, SNAI2, CCL4 and DKC1 (Fig. 1C, D).

**Figure 1.**
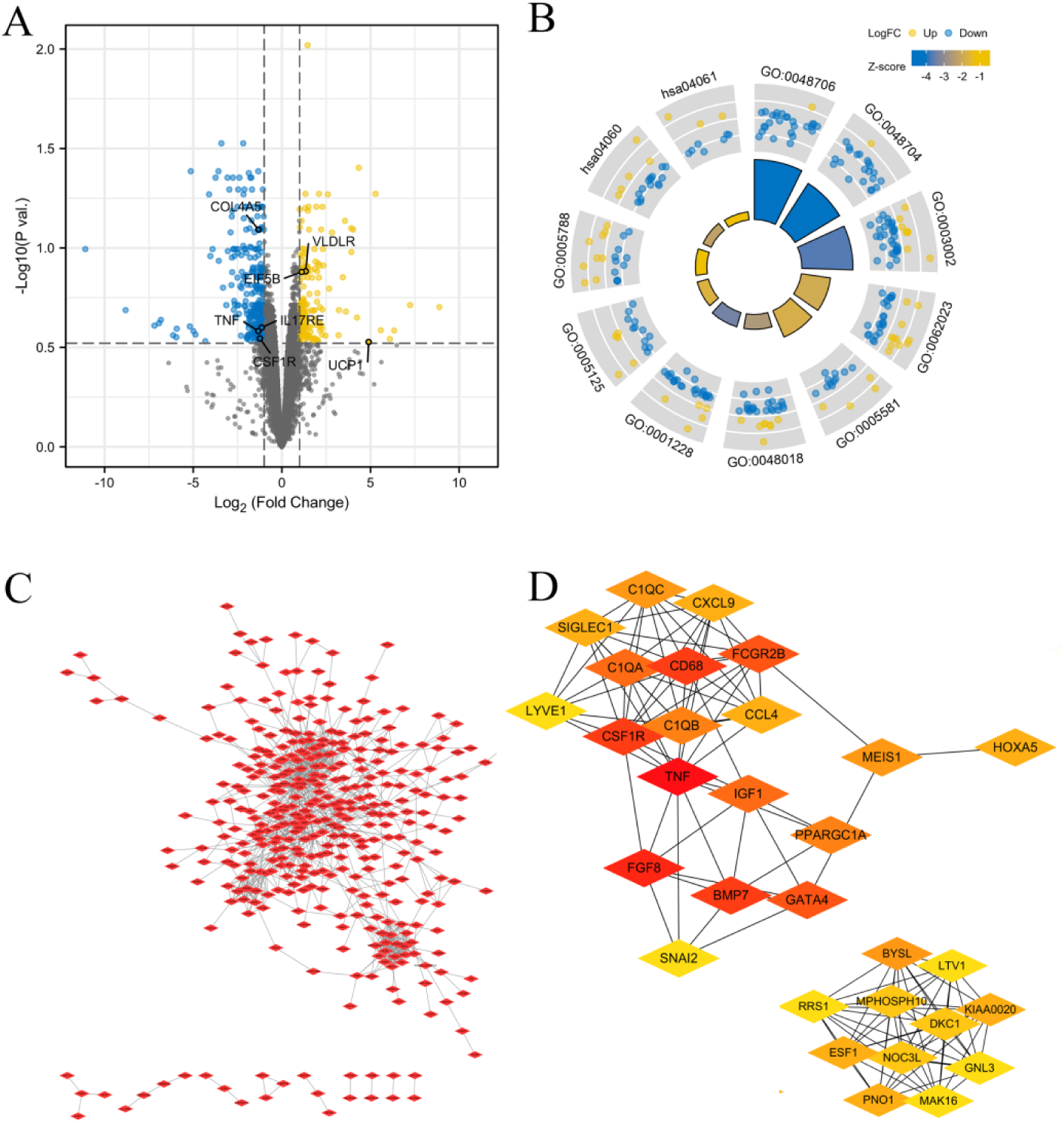
DEGs between EAT from patients with AF and subjects with SR. (A) The volcano plot of DEGs between EAT from patients with AF and subjects with SR. (B) The circle diagram of the GO/KEGG pathways enriched by DEGs between EAT from patients with AF and subjects with SR. (C-D) The PPI network (C) and the top30-degree hub genes (D) were obtained between EAT from patients with AF and subjects with SR.

**Table 1.**
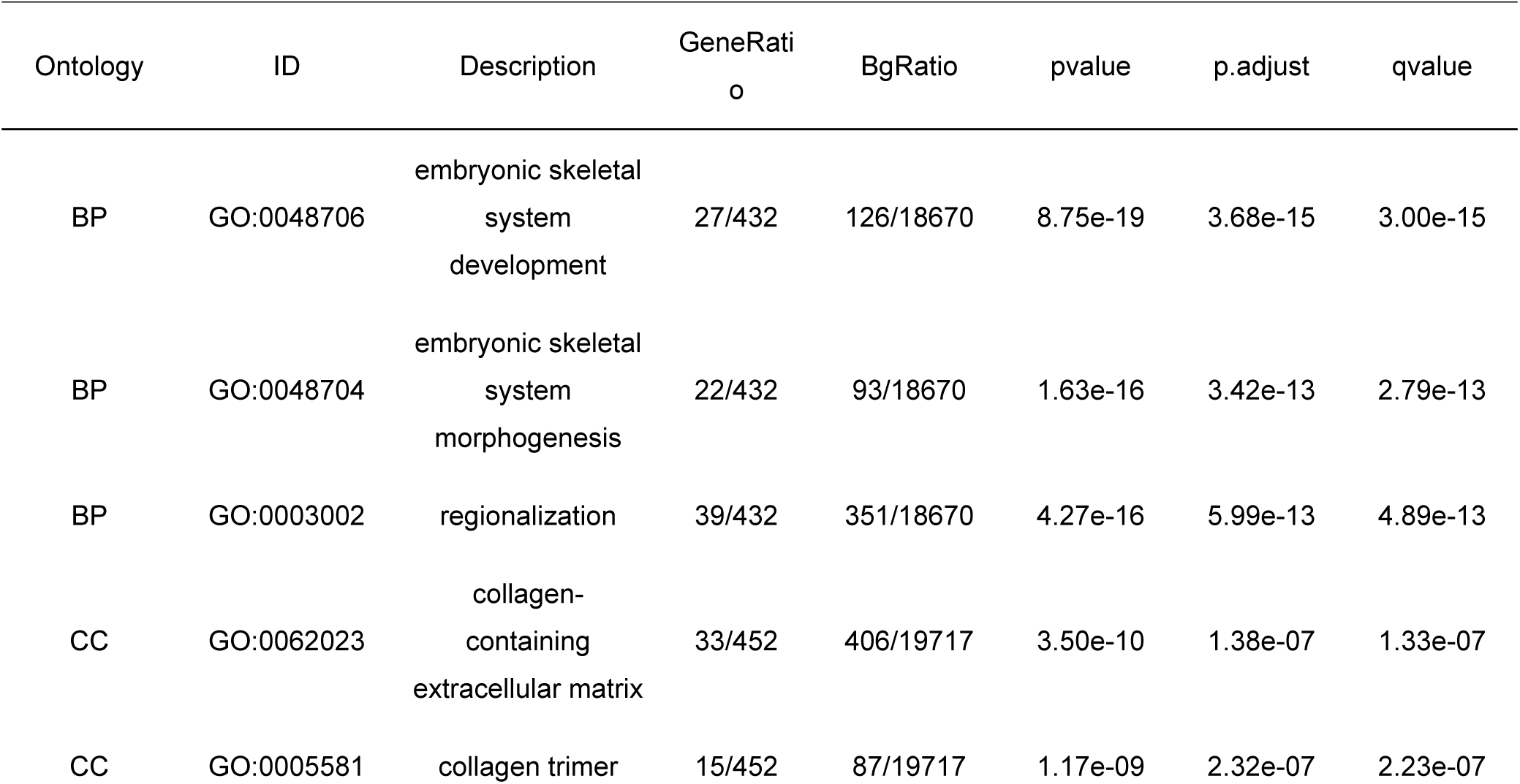
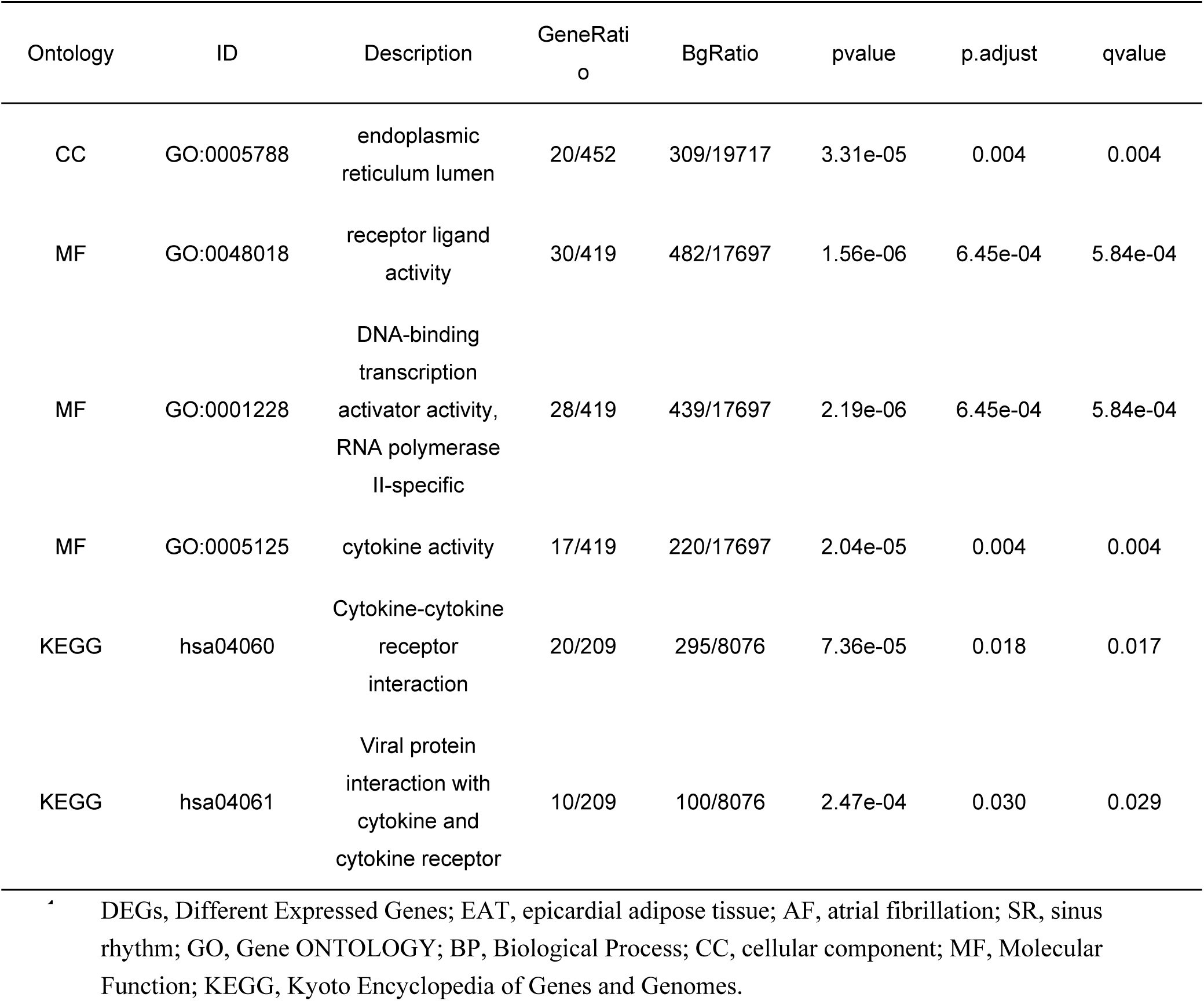
The GO/KEGG pathways enriched by DEGs between EAT from subjects with AF and SR.

| Ontology | ID | Description | GeneRatio | BgRatio | pvalue | p.adjust | qvalue |
| --- | --- | --- | --- | --- | --- | --- | --- |
| BP | GO:0048706 | embryonic skeletal system development | 27/432 | 126/18670 | 8.75e-19 | 3.68e-15 | 3.00e-15 |
| BP | GO:0048704 | embryonic skeletal system morphogenesis | 22/432 | 93/18670 | 1.63e-16 | 3.42e-13 | 2.79e-13 |
| BP | GO:0003002 | regionalization | 39/432 | 351/18670 | 4.27e-16 | 5.99e-13 | 4.89e-13 |
| CC | GO:0062023 | collagen-containing extracellular matrix | 33/452 | 406/19717 | 3.50e-10 | 1.38e-07 | 1.33e-07 |
| CC | GO:0005581 | collagen trimer | 15/452 | 87/19717 | 1.17e-09 | 2.32e-07 | 2.23e-07 |
| CC | GO:0005788 | endoplasmic reticulum lumen | 20/452 | 309/19717 | 3.31e-05 | 0.004 | 0.004 |
| MF | GO:0048018 | receptor ligand activity | 30/419 | 482/17697 | 1.56e-06 | 6.45e-04 | 5.84e-04 |
| MF | GO:0001228 | DNA-binding transcription activator activity, RNA polymerase II-specific | 28/419 | 439/17697 | 2.19e-06 | 6.45e-04 | 5.84e-04 |
| MF | GO:0005125 | cytokine activity | 17/419 | 220/17697 | 2.04e-05 | 0.004 | 0.004 |
| KEGG | hsa04060 | Cytokine-cytokine receptor interaction | 20/209 | 295/8076 | 7.36e-05 | 0.018 | 0.017 |
| KEGG | hsa04061 | Viral protein interaction with cytokine and cytokine receptor | 10/209 | 100/8076 | 2.47e-04 | 0.030 | 0.029 |

DEGs, Different Expressed Genes; EAT, epicardial adipose tissue; AF, atrial fibrillation; SR, sinus rhythm; GO, Gene ONTOLOGY; BP, Biological Process; CC, cellular component; MF, Molecular Function; KEGG, Kyoto Encyclopedia of Genes and Genomes.

### 3.2 Identification of DEGs of PAT and further analysis

A total of 190 up-regulated and 149 down-regulated DEGs was obtained between whole PAT and SAT, which were mainly involved in leukocyte migration, cellular response to chemokine, immunological synapse, Cytokine-cytokine receptor interaction, and heparin binding. A total of 115 up-regulated and 53 down-regulated DEGs was obtained between isolated PAT and SAT, which were mainly involved in NF-kappa B signaling pathway, membrane raft, Complement and coagulation cascades, and regulation of lymphocyte activation (Fig. 2A-C; Tab. S1, S2). Utilizing STRING and Cytoscape for isolated PAT, the top15-degree hub genes were obtained, including PAX5, CCL4, CR2, WNT5A, CD8A and SELL (Fig. 2D, E). Using immune infiltration analysis in whole PAT related DEGs, the abundance of M0 macrophages was significantly different between PAT and SAT and 22 M0 macrophages-related DEGs were also obtained, such as MMP7, APOE, CCND1, MMP12 and APOC1 (Fig. 2F, G; Tab. S3).

**Figure 2.**
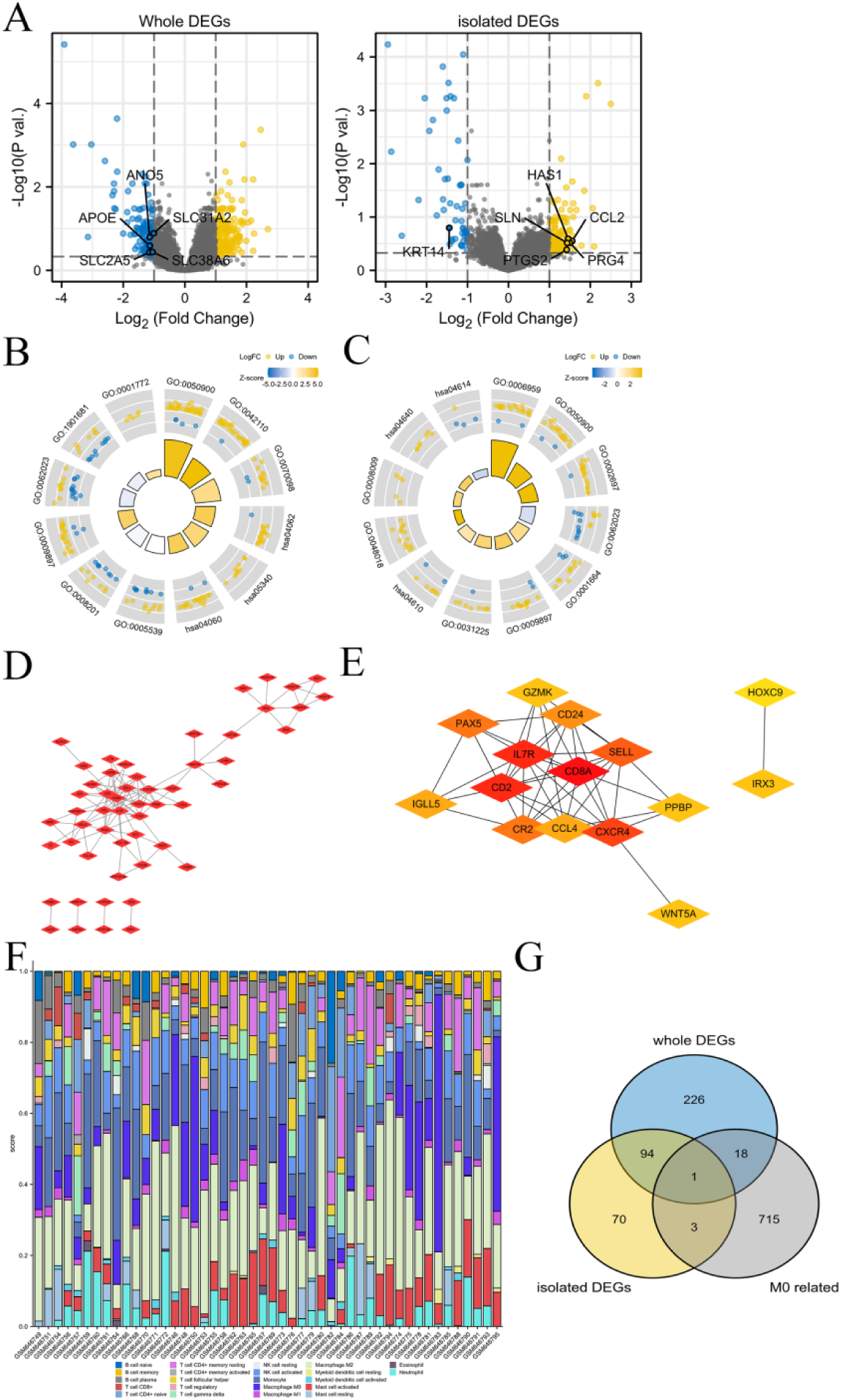
DEGs between PAT and SAT. (A) The volcano plot of DEGs between whole PAT or isolated PAT and SAT. (B-C) The bar plot of whole PAT-related DEGs (B) and isolated PAT-related DEGs (C). (D-E) The PPI network (D) and the hub genes (E) were obtained from isolated PAT-related DEGs. (G) The immune cells abundance in whole PAT using Cibersort. (H) Venn diagram of M0 macrophages-related DEGs in whole PAT or isolated PAT.

### 3.3 Senescence-related DEGs and further analysis

The senescence-related DEGs in EAT were obtained from GSE35957, which were mainly involved in mitotic nuclear division, chromosomal region, PI3K-Akt signaling pathway, and Cell cycle (Fig. 3A, B; Tab. 2). Venn diagram was utilized to obtain the same transcripts and PPI network was used to 94 overlapping genes. The top15-degree hub genes were explored, including BMP7, COL4A3, COL4A4, CHRD, IRX1, SPON1, ALPL, CTSK, TCF19 and SIX2 (Fig. 3C-E). The PPI network of non-senescence DEGs was constructed and the hub genes were also obtained, including CD68, C1QC, CSF1R, CCL4 and GATA4 (Fig. S2).

**Figure 3.**
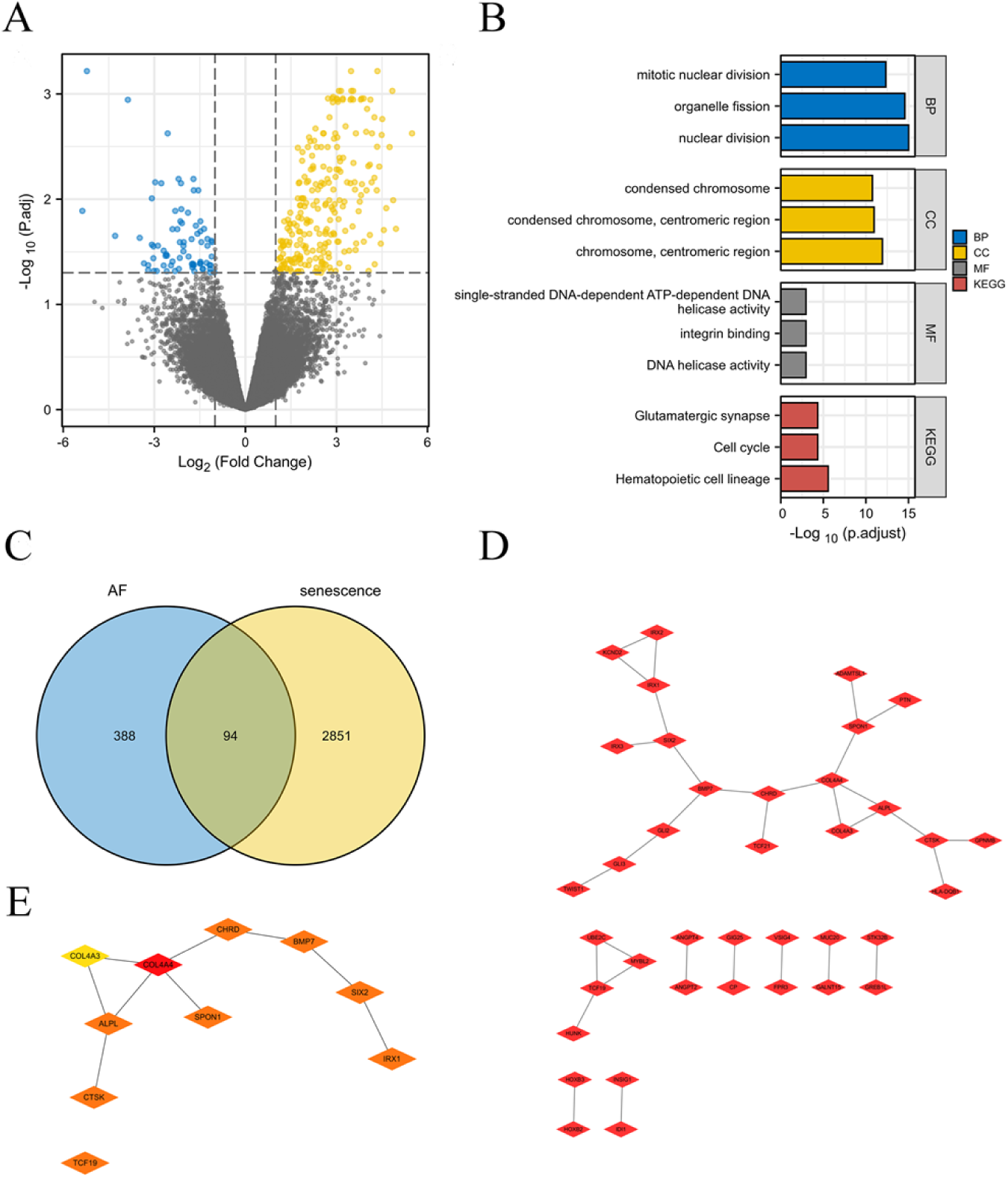
The senescence-related DEGs in EAT from patients with AF. (A) The volcano plot of the senescence-related DEGs in MSCs. (B) The GO/KEGG pathways enriched by the senescence-related DEGs in MSCs. (C) Venn diagram of the senescence-related DEGs and DEGs between EAT from patients with AF and subjects with SR. (D-E) The PPI network (E) and the 10 hub genes (F) were obtained from the screened 94 genes.

**Table 2.** The GO/KEGG pathways enriched by senescence-related DEGs in MSCs.

| ONTOLOGY | ID | Description | GeneRatio | BgRatio | pvalue | p.adjust | qvalue |
| --- | --- | --- | --- | --- | --- | --- | --- |
| BP | GO:0000280 | nuclear division | 112/2130 | 407/18670 | 1.59e-19 | 9.70e-16 | 8.01e-16 |
| BP | GO:0048285 | organelle fission | 118/2130 | 449/18670 | 8.87e-19 | 2.71e-15 | 2.23e-15 |
| BP | GO:0140014 | mitotic nuclear division | 79/2130 | 264/18670 | 2.27e-16 | 4.62e-13 | 3.81e-13 |
| BP | GO:0007059 | chromosome segregation | 86/2130 | 321/18670 | 1.51e-14 | 2.31e-11 | 1.90e-11 |
| BP | GO:0098813 | nuclear chromosome segregation | 73/2130 | 262/18670 | 1.78e-13 | 1.90e-10 | 1.57e-10 |
| CC | GO:0000775 | chromosome, centromeric region | 63/2227 | 193/19717 | 1.72e-15 | 1.18e-12 | 1.01e-12 |
| CC | GO:0000779 | condensed chromosome, centromeric region | 45/2227 | 118/19717 | 3.24e-14 | 1.11e-11 | 9.56e-12 |
| CC | GO:0000793 | condensed chromosome | 66/2227 | 223/19717 | 7.57e-14 | 1.73e-11 | 1.49e-11 |
| CC | GO:0000776 | kinetochore | 48/2227 | 135/19717 | 1.04e-13 | 1.78e-11 | 1.53e-11 |
| CC | GO:0098687 | chromosomal region | 87/2227 | 349/19717 | 5.25e-13 | 7.22e-11 | 6.20e-11 |
| MF | GO:0003678 | DNA helicase activity | 26/2130 | 81/17697 | 1.56e-06 | 0.001 | 0.001 |
| MF | GO:0005178 | integrin binding | 35/2130 | 132/17697 | 4.15e-06 | 0.001 | 0.001 |
| MF | GO:0017116 | single-stranded DNA-dependent ATP-dependent DNA helicase activity | 11/2130 | 20/17697 | 4.44e-06 | 0.001 | 0.001 |
| MF | GO:0043142 | single-stranded DNA-dependent ATPase activity | 11/2130 | 20/17697 | 4.44e-06 | 0.001 | 0.001 |
| MF | GO:0035173 | histone kinase activity | 10/2130 | 17/17697 | 5.44e-06 | 0.001 | 0.001 |
| KEGG | hsa04640 | Hematopoietic cell lineage | 34/992 | 99/8076 | 8.53e-09 | 2.73e-06 | 2.26e-06 |
| KEGG | hsa04110 | Cell cycle | 36/992 | 124/8076 | 4.07e-07 | 4.53e-05 | 3.76e-05 |
| KEGG | hsa04724 | Glutamatergic synapse | 34/992 | 114/8076 | 4.25e-07 | 4.53e-05 | 3.76e-05 |
| KEGG | hsa04015 | Rap1 signaling pathway | 49/992 | 210/8076 | 4.88e-06 | 3.90e-04 | 3.24e-04 |
| KEGG | hsa04151 | PI3K-Akt signaling pathway | 72/992 | 354/8076 | 7.31e-06 | 4.68e-04 | 3.88e-04 |

1 DEGs, Different Expressed Genes; MSCs, Mesenchymal stem cells; GO, Gene ONTOLOGY; BP, 2 Biological Process; CC, cellular component; MF, Molecular Function; KEGG, Kyoto Encyclopedia 3 of Genes and Genomes.

### 3.4 The hub genes and their interactions

The screened DEGs and their interactions were demonstrated (Fig. 4). TFs, miRNA, drugs, and chemicals networks interactions with the hub DEGs were constructed, which can further provide the insight for treatment to aging patients with AF.

**Figure 4.**
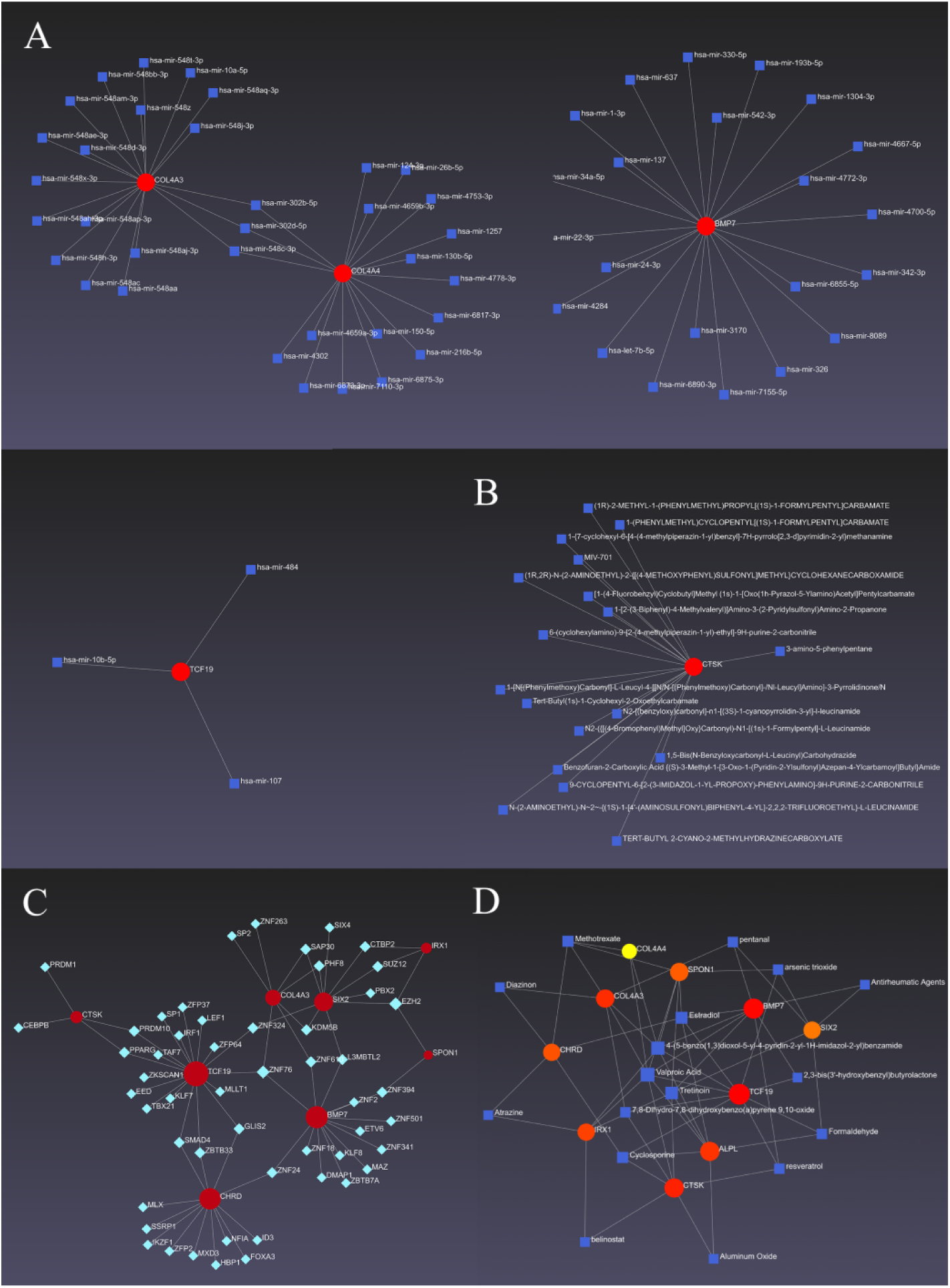
The hub genes and their interactions. The networks of miRNA-the hub DEGs (A), drugs-the hub DEGs (B), TFs-the hub DEGs (C) and chemicals-the hub DEGs (D) were constructed.

### 3.5 The central-marginal dispersion in EAT from patients with AF

To explore the central-marginal dispersion in EAT from patients with AF, GSE154436 was downloaded and analyzed to obtain DEGs, which were mainly involved in extracellular matrix organization, collagen-containing extracellular matrix, and ATPase binding (Fig. S3A, B; Tab. S4). Venn diagram was utilized to obtain the same transcripts between AF-related DEGs and central-marginal dispersion-related DEGs and 6 marginal-related DEGs were screened, such as C7 and HOXB3, and 1 central-related DEG (CYSLTR2) was explored (Fig. S3C).

### 3.6 Human samples Validation

To investigate senescence-related hub genes effects on AF progression, We selected 50 cases of persistent atrial fibrillation and 50 cases of paroxysmal atrial fibrillation that were diagnosed and hospitalized in the Department of Cardiology of the First Affiliated Hospital of Hebei North University from February 1st 2025 to October 30th 2025. There were no significant differences at baseline characteristics and laboratory examinations at 1d after admission except heart failure history and function examinations, for instance, ALT, LDL, LVEF and LA diameters (Tab. S5). Using univariate analysis and multivariate analysis, heart failure history, ALT and LA diameters can be prognostic factors for AF progression (Tab. S6).

BMP7, COL4A3, COL4A4, CHRD, IRX1, SPON1, ALPL, TCF19 and SIX2 were validated to be highly expressed in persistent atrial fibrillation compared to paroxysmal atrial fibrillation, which may be potential therapeutic targets to treat AF progression and improve patients’ prognosis (Fig. 5).

**Figure 5.**
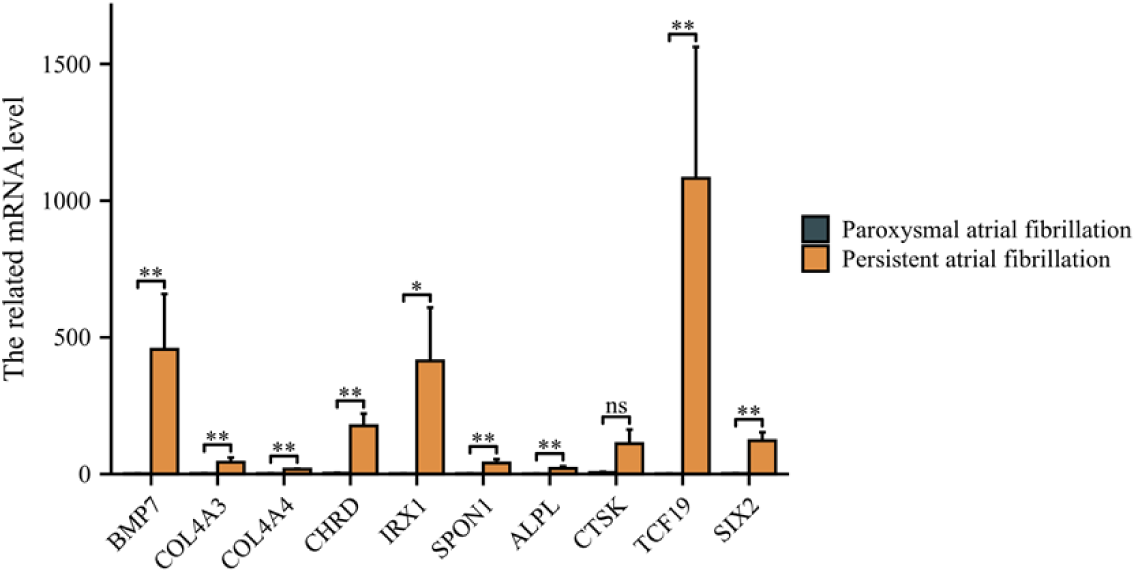
The hub genes were validated using human blood samples. qPCR was performed in human blood tissues between with persistent atrial fibrillation and paroxysmal atrial fibrillation. * P<0.05; **P<0.01; ns, not significant.

## 4. Discussion

Atrial Fibrillation (AF) is one of the most common types of arrhythmia, with its incidence significantly increasing with age. AF is associated with a variety of potential cardiovascular and metabolic diseases, and it is linked to increased morbidity and mortality. As the population ages, AF has an increasingly significant impact on public health and economic burden[10]. In 2020, AF affected 37.5 million patients worldwide, approximately 0.51% of the global population. The prevalence of AF is 1–2% which rises to around 10% by the time we reach 65 years of age[8].

According to research by Samer Asmar and colleagues, the prevalence of AF in the United States is close to 4 million patients, and it is expected that the incidence will double in the next 50 years. The hospitalization costs for AF patients are three times those of patients without AF, adding 6 billion to 26 billion in medical expenses each year[11]. A review by E.-H. Diallo and others studied the predictive factors and impact of AF after thoracic surgery, involving 39,783 patients, and found that the average hospital stay for AF patients was extended by 3 days[12]. AF is a disease closely related to ageing, and its epidemiology reveals that with the ageing of the population, the incidence of AF and the associated medical burden will continue to increase. In addition, early detection and treatment of AF are crucial for preventing complications, and wearable devices provide a new means of detection that can help identify undiagnosed AF patients on a large scale[13].

Cellular senescence is generally an irreversible proliferative arrest in damaged normal cells that have exited the cell cycle[14]. In 1961, Hayflick and Moorehead first studied the permanent arrest of proliferation at the cellular level. They found that the lifespan of primary human cells was limited to approximately 60 cell divisions[15]. Cellular senescence can be divided into three main categories: (i) replicative senescence; (ii) developmentally programmed cellular senescence, evidenced to be a crucial process for healthy embryonic development; and (iii) stress-induced premature senescence, triggered by a wide range of external and internal by stimuli such as oxidative stress, oncogene expression, DNA damage[16].

CHRD (Chromosome Healing and Repair Detector) is a chromatin remodeling factor gene. The protein encoded by this gene can affect gene expression and chromosome stability by regulating the structure of chromatin. By protecting telomeres and repairing DNA damage, it helps maintain genome stability and is closely related to cell ageing[17]. CHRDL1 (chordin-like 1), a new type of fat hormone, can significantly promote lipogenesis and lipid accumulation[18].

Bone Morphogenetic Protein 7 (BMP7), namely bone morphogenetic protein 7 gene, belongs to a member of the transforming growth factor-β (TGF-β) super-family[19]. The BMP7 gene encodes a protein that has the characteristics of a secreted glycoprotein. In the body, BMP7 activates signal transduction pathways by binding to specific receptors on the cell surface, thereby affecting cell fate and function[20]. The BMP7 signaling pathway plays a role in several biological processes, such as neuronal differentiation, heart development, liver regeneration, and tumorigenesis.

COL4A3 (Collagen, type IV, alpha 3) and COL4A4 (Collagen, type IV, alpha 4) are both genes encoding collagen type IV. Collagen IV is an important basement membrane protein, which is mainly found in the extracellular matrix of organs such as the kidneys, ears and eyes, and plays a role in supporting and protecting cells. The genetic mutations of the two types of collagen may lead to changes in the structure and function of type IV collagen, thus affecting the normal function of body organs and causing some autosomal recessive genetic diseases[21,22]. APP gene (Amyloid Precursor Protein), is a coding gene located on human chromosome 21. The APP gene is mainly expressed in tissues such as the human liver, gut and brain, and the Protein it encodes is called Amyloid Precursor Protein (APP). As a glycoprotein, APP plays an important role in maintaining neuronal homeostasis, such as signaling, neuronal development, and intracellular transport[23]. Mutations or abnormal expression of APP genes can lead to the occurrence of many diseases such as Alzheimer’s disease[24].

There are some limitations. The sample size of human validation is quite small. The large, multi-center and double-blind cohort is still required to investigate the cell senescence effects on AF.

## 5. Conclusion

Based on our current study, our research provided bioinformatics analysis about the prognostic targets for patients with AF. The screened hub genes, adipocyte-related senescent genes, were validated using GEO dataset and human blood samples, which may be therapeutic targets AF patients to prevent myocardiopathy progression and adverse cardiovascular events.

## Declarations

### Ethics approval and consent to participate

Before participating in the study, all individuals provided written informed consent, and the protocols were approved by the First Affiliated Hospital of Hebei North University[K2023059].

### Author contributions

Conceptualization: Feixing Li, Fangjiang Li.

Data curation: Feixing Li, Linbing Liu, Xiaoyuan Wang.

Formal analysis:Feixing Li, Wei Liu, Meiling Du.

Investigation: Mengyang Yi Yue Shi.

Methodology: Feixing Li, Aiai Zhang.

Software:Feixing Li, Wei Han.

Supervision: Feixing Li, Jianying Song.

Writing – original draft: Feixing Li.

Writing – review & editing: Feixing Li.

### Consent for publication

All the authors have agreed to the submission and publication of this paper.

### Availability of data and materials

Not applicable.

### Competing interests

On behalf of all authors, the corresponding author stated that there is no conflict of interest and agreed to submit it.

### Funding

This work was funded by the Sci-tech Plan Project of Zhangjiakou (2221156D) and Medical Science Research Plan of Hebei Province(20242068,20242338).

## Acknowledgement

We would like to express our gratitude to all those who helped us during the writing of this manuscript. Thanks to all the peer reviewers for their opinions and suggestions.

## Supplementary Figure Legend

Supplementary Figure 1. The bar plot (A) and PCA diagram (B) between EAT from patients with AF and subjects with SR.

Supplementary Figure 2. The PPI network (A) and the hub genes (B) were explored from non-senescence DEGs in EAT from patients with AF.

Supplementary Figure 3. The central-marginal dispersion in EAT from patients with AF. (A) The volcano plot of the central-marginal dispersion-related DEGs. (B) The circle diagram of GO/KEGG pathways enriched by the central-marginal dispersion-related DEGs. (C) Venn diagram was utilized to obtain the same transcripts between AF-related DEGs and central-marginal dispersion-related DEGs. 6 marginal-related DEGs and 1 central-related DEG was screened.

## Notes

### Competing Interest Statement

The authors have declared no competing interest.

## Reference

[1] Assaf A, Mekhael M, Noujaim C, et al. Effect of fibrosis regionality on atrial fibrillation recurrence: insights from DECAAF II. Eur Eur Pacing Arrhythm Card Electrophysiol J Work Groups Card Pacing Arrhythm Card Cell Electrophysiol Eur Soc Cardiol. 2023;25(9):euad199. doi:10.1093/europace/euad199.

[2] Hopman LHGA, Visch JE, Bhagirath P, et al. Right atrial function and fibrosis in relation to successful atrial fibrillation ablation. Eur Heart J Cardiovasc Imaging. 2023;24(3):336–345. doi:10.1093/ehjci/jeac152.

[3] Kim J, Park SJ, Jeong DS, et al. Left atrial strain predicts fibrosis of left atrial appendage in patients with atrial fibrillation undergoing totally thoracoscopic ablation. Front Cardiovasc Med. 2023;10:1130372. doi:10.3389/fcvm.2023.1130372.

[4] Chahine Y, Askari-Atapour B, Kwan KT, et al. Epicardial adipose tissue is associated with left atrial volume and fibrosis in patients with atrial fibrillation. Front Cardiovasc Med. 2022;9:1045730. doi:10.3389/fcvm.2022.1045730.

[5] Nezami Z, Holm H, Ohlsson M, et al. The impact of myocardial fibrosis biomarkers in a heart failure population with atrial fibrillation-The HARVEST-Malmö study. Front Cardiovasc Med. 2022;9:982871. doi:10.3389/fcvm.2022.982871.

[6] Ma S, Yan F, Hou Y. Intermedin 1-53 Ameliorates Atrial Fibrosis and Reduces Inducibility of Atrial Fibrillation via TGF-β1/pSmad3 and Nox4 Pathway in a Rat Model of Heart Failure. J Clin Med. 2023;12(4):1537. doi:10.3390/jcm12041537.

[7] Adili A, Zhu X, Cao H, et al. Atrial Fibrillation Underlies Cardiomyocyte Senescence and Contributes to Deleterious Atrial Remodeling during Disease Progression. Aging Dis. 2022;13(1):298–312. doi:10.14336/AD.2021.0619.

[8] Guo G, Watterson S, Zhang SD, et al. The role of senescence in the pathogenesis of atrial fibrillation: A target process for health improvement and drug development. Ageing Res Rev. 2021;69:101363. doi:10.1016/j.arr.2021.101363.

[9] Jesel L, Abbas M, Park SH, et al. Atrial Fibrillation Progression Is Associated with Cell Senescence Burden as Determined by p53 and p16 Expression. J Clin Med. 2019;9(1):36. doi:10.3390/jcm9010036.

[10] Bizhanov KA, Аbzaliyev KB, Baimbetov AK, Sarsenbayeva AB, Lyan E. Atrial fibrillation: Epidemiology, pathophysiology, and clinical complications (literature review). J Cardiovasc Electrophysiol. 2023;34(1):153–165. doi:10.1111/jce.15759.

[11] Chyou JY, Barkoudah E, Dukes JW, et al. Atrial Fibrillation Occurring During Acute Hospitalization: A Scientific Statement From the American Heart Association. Circulation. 2023;147(15):e676–e698. doi:10.1161/CIR.0000000000001133

[12] Diallo EH, Brouillard P, Raymond JM, Liberman M, Duceppe E, Potter BJ. Predictors and impact of postoperative atrial fibrillation following thoracic surgery: a state-of-the-art review. Anaesthesia. 2023;78(4):491–500. doi:10.1111/anae.15957.

[13] Lubitz SA, Faranesh AZ, Selvaggi C, et al. Detection of Atrial Fibrillation in a Large Population Using Wearable Devices: The Fitbit Heart Study. Circulation. 2022;146(19):1415–1424. doi:10.1161/CIRCULATIONAHA.122.060291.

[14] Roger L, Tomas F, Gire V. Mechanisms and Regulation of Cellular Senescence. Int J Mol Sci. 2021 Dec 6;22(23):13173. doi: 10.3390/ijms222313173.

[15] Hayflick, L.; Moorhead, P.S. The Serial Cultivation of Human Diploid Cell Strains. Exp. Cell Res. 1961, 25, 585–621. [

[16] Lucas V, Cavadas C, Aveleira CA. Cellular Senescence: From Mechanisms to Current Biomarkers and Senotherapies. Pharmacol Rev. 2023 Jul;75(4):675–713. doi: 10.1124/pharmrev.122.000622.

[17] Zhou Y, Cao G, Guan Z, Mao C. Chordin-Like 2: A Possible Therapeutic Target for Gastric Cancer by Affecting Cell Cycle and Proliferation. J Oncol. 2022 Nov 8;2022:4607715. doi: 10.1155/2022/4607715.

[18] Ahn J, Suh Y, Lee K. Chordin-like 1, a Novel Adipokine, Markedly Promotes Adipogenesis and Lipid Accumulation. Cells. 2023 Feb 15;12(4):624. doi: 10.3390/cells12040624.

[19] Saini S, Duraisamy AJ, Bayen S, Vats P, Singh SB. Role of BMP7 in appetite regulation, adipogenesis, and energy expenditure. Endocrine. 2015 Mar;48(2):405–9. doi: 10.1007/s12020-014-0406-8.

[20] Ripmeester EGJ, Caron MMJ, van den Akker GGH, Steijns J, Surtel DAM, Cremers A, Peeters LCW, van Rhijn LW, Welting TJM. BMP7 reduces the fibrocartilage chondrocyte phenotype. Sci Rep. 2021 Oct 4;11(1):19663. doi: 10.1038/s41598-021-99096-0.

[21] Xia L, Cao Y, Guo Y, et al. A Novel Heterozygous Mutation of the COL4A3 Gene Causes a Peculiar Phenotype without Hematuria and Renal Function Impairment in a Chinese Family. Dis Markers. 2019 Feb 10;2019:8705989. doi: 10.1155/2019/8705989.

[22] Fan LL, Liu L, Luo FM, et al. A novel heterozygous variant of the COL4A4 gene in a Chinese family with hematuria and proteinuria leads to focal segmental glomerulosclerosis and chronic kidney disease. Mol Genet Genomic Med. 2020 Dec;8(12):e1545. doi: 10.1002/mgg3.154.

[23] Li R, Song J, Zhao A, Diao X, et al. Association of APP gene polymorphisms and promoter methylation with essential hypertension in Guizhou: a case-control study. Hum Genomics. 2023 Mar 20;17(1):25. doi: 10.1186/s40246-023-00462-y.

[24] Reddy PH, Oliver DM. Amyloid Beta and Phosphorylated Tau-Induced Defective Autophagy and Mitophagy in Alzheimer’s Disease. Cells. 2019 May 22;8(5):488. doi: 10.3390/cells8050488.

